# Degraded myelin is associated with cold hypersensitivity in paclitaxel-induced peripheral neuropathy

**DOI:** 10.64898/2026.05.29.728780

**Authors:** Laura R Osborn, Mary Grace Bishop, Kayleigh A Rodriguez, Dakota M Redling, Renee Fox, Ashley M Ezeldin, Emma I Zheng, Kimberly E Stephens

## Abstract

Cold allodynia is a debilitating symptom of chemotherapy-induced peripheral neuropathy (CIPN), a side effect of many chemotherapeutics, including paclitaxel (PTX). CIPN is highly prevalent and can lead to cessation of chemotherapy, threatening cancer patient survival. PTX induces cold-evoked pain in 60% of CIPN patients and this pain is a predictor for more severe neuropathy. Cold is transduced by primary sensory neurons in dorsal root ganglia (DRG). PTX can induce maladaptive changes in gene expression that lead to dysregulated nociceptor function. Previous work examining PTX-induced transcriptional changes have been unable to isolate mechanisms associated with pathological cold pain. The epigenetic changes that underlie long-lasting transcriptional changes in PTX-induced persistent cold hypersensitivity are not known.

Following three consecutive cycles of PTX, we find that mice continue to exhibit cold hypersensitivity for a minimum of four weeks following the resolution of PTX-induced mechanical hypersensitivity. This exclusive cold hypersensitivity phenotype allows us to examine the mechanisms that specifically underlie pathological cold pain. During this period of PTX-induced cold hypersensitivity, we found decreased expression of genes whose expression is necessary for myelin sheath formation and maintenance, degraded myelin sheaths in the distal sciatic nerve, and decreased chromatin accessibility at genomic regions that contain binding motifs for pro-myelinating transcription factors.

Our findings suggest that epigenetic-regulated myelin programs perpetuate myelin degradation and drive pathological cold pain. Our improved understanding of the specific mechanisms that maintain myelin degradation during cold hypersensitivity will improve cold pain management, facilitate the completion of cancer treatment, and improve the quality of life for chronic pain patients.

## Introduction

Cold allodynia is a particularly debilitating symptom of chemotherapy-induced peripheral neuropathy (CIPN) and other neuropathic pain conditions, such as multiple sclerosis and fibromyalgia, where patients experience reduced cold pain thresholds and increased cold pain intensity.^1–3^ CIPN is a dose-dependent side effect of many chemotherapeutics and affects more than 60% of the 1 million cancer survivors in the US annually.^4–6^ Current treatments for CIPN, such as duloxetine, calcium channel modulators, cannabinoids, and opioids, have limited efficacy, are associated with poorly tolerated side effects, and do not specifically target cold pain.^7–9^ The debilitating effects of cold allodynia on patients underscores the importance of investigating the mechanisms that underlie cold pain to support the development of targeted and effective therapies.

Paclitaxel (PTX) is a commonly used chemotherapeutic that damages peripheral nerves, induces sensory abnormalities, and produces painful neuropathies in up to 97% of patients.^10–12^ The intensity of PTX-induced peripheral neuropathy (PIPN) increases with cumulative chemotherapy dose and can last up to 11 years after chemotherapy is completed.^5,11,13,14^ Cold allodynia can be irreversible between chemotherapy cycles, and experiencing cold pain after early cycles of chemotherapy is a predictor for the development of chronic and severe neuropathy.^15^ During cancer therapy, PIPN development can lead to dose-reduction or even cessation of chemotherapy which threatens the cancer patient’s survival.^11,12,16^

During CIPN, changes in the expression of ion channels and various other factors in the dorsal root ganglia (DRG) promote increased neuronal excitability and pain development.^10,17–19^ Epigenetic mechanisms regulate transcription by altering chromatin structure to promote adaptation of the cells in response to intrinsic and environmental stimuli. However, some epigenetic changes are associated with maladaptive sensory neuron function and the development of hypersensitivity in rodent models of CIPN.^20–23^ Cold hypersensitivity has been associated with changes in *Trpm8* and potassium channel expressions.^24–32^ Despite our advances in understanding the mechanisms of cold hypersensitivity, efficacious treatments for pathological cold pain have not been developed.^7–9^ However, previous work examining transcriptional changes during cold hypersensitivity have been unable to disaggregate findings due to the presence of confounding mechanical hypersensitivity. The persistent changes in gene transcription that underlie exclusive cold hypersensitivity are not known.

Using a translational PIPN model, we observe a period of exclusive cold hypersensitivity that lasts at least four weeks after mechanical hypersensitivity returns to baseline and, therefore, provides a unique opportunity to examine the mechanisms underlying pathological cold pain.^33^ In this study, we examine the transcriptome and genome-wide chromatin accessibility profiles in the DRG to identify the mechanisms associated with chronic cold hypersensitivity.

## Methods

### Animals

Adult female (8 weeks old) C57BL/6J mice were obtained from Jackson Laboratories (Bar Harbor, ME). Mice were housed in a room with controlled temperature (22±1°C) and humidity (60±10%), a 12-hour light/dark cycle, and *ad libitum* access to standard rodent chow and water. Mice acclimated to the housing facility for >48 hours prior to experimental procedures and all procedures were performed during the animal’s light cycle (7:00am to 7:00pm). Mice were randomly assigned to a treatment group (n=12/group) upon arrival and all mice in a cage received the same treatment. Behavioral assessments were conducted on eight mice from each group and the remaining four mice per group were used for tissue collection. All procedures involving animals were reviewed and approved by the University of Arkansas for Medical Sciences Institutional Animal Care and Use Committee (IACUC) and were performed in accordance with the National Institutes of Health’s Guide for the Care and Use of Laboratory Animals.

### Paclitaxel

Paclitaxel (Athenex, Schaumburg, IL) was obtained from Arkansas Children’s Hospital Pharmacy as 6 mg/kg paclitaxel dissolved in 527 mg Polyoxyl 35 Castor Oil NF, 49.7% (V/V) United States Pharmacopeia Grade Dehydrated Alcohol, and 2 mg Citric Acid. PTX was diluted to 0.5 mg/mL in sterile 0.9% sodium chloride immediately prior to use. Control mice received vehicle (VEH) only [5% ethanol, 5% Cremophor EL (Merck KGaA, Darmstadt, Germany), 95% sterile 0.9% sodium chloride].

### Experimental design

Adult female (8 weeks old) C57BL/6J mice were randomly assigned to either PTX (n=12) or VEH (n=12) groups. Baseline behavioral assessments were conducted prior to administration of the first cycle. Each cycle consisted of four intraperitoneal (IP) injections of PTX (4 mg/kg) or vehicle, with injections given every other day for a dosage of 16 mg/kg/cycle. Each animal received a total of three cycles (cumulative dosage of 48 mg/kg) as previously described (Fig. 1A).^33–35^ Subsequent cycles of PTX or VEH were given once mechanical sensitivity returned to baseline in the behavior group following the previous cycle (i.e. 28-30 days after the first dose of cycles 1 and 2) (Fig. 1A). Consecutive cycles of PTX and VEH did not have noticeable impacts on rodent overall health.^33^

**Figure 1.**
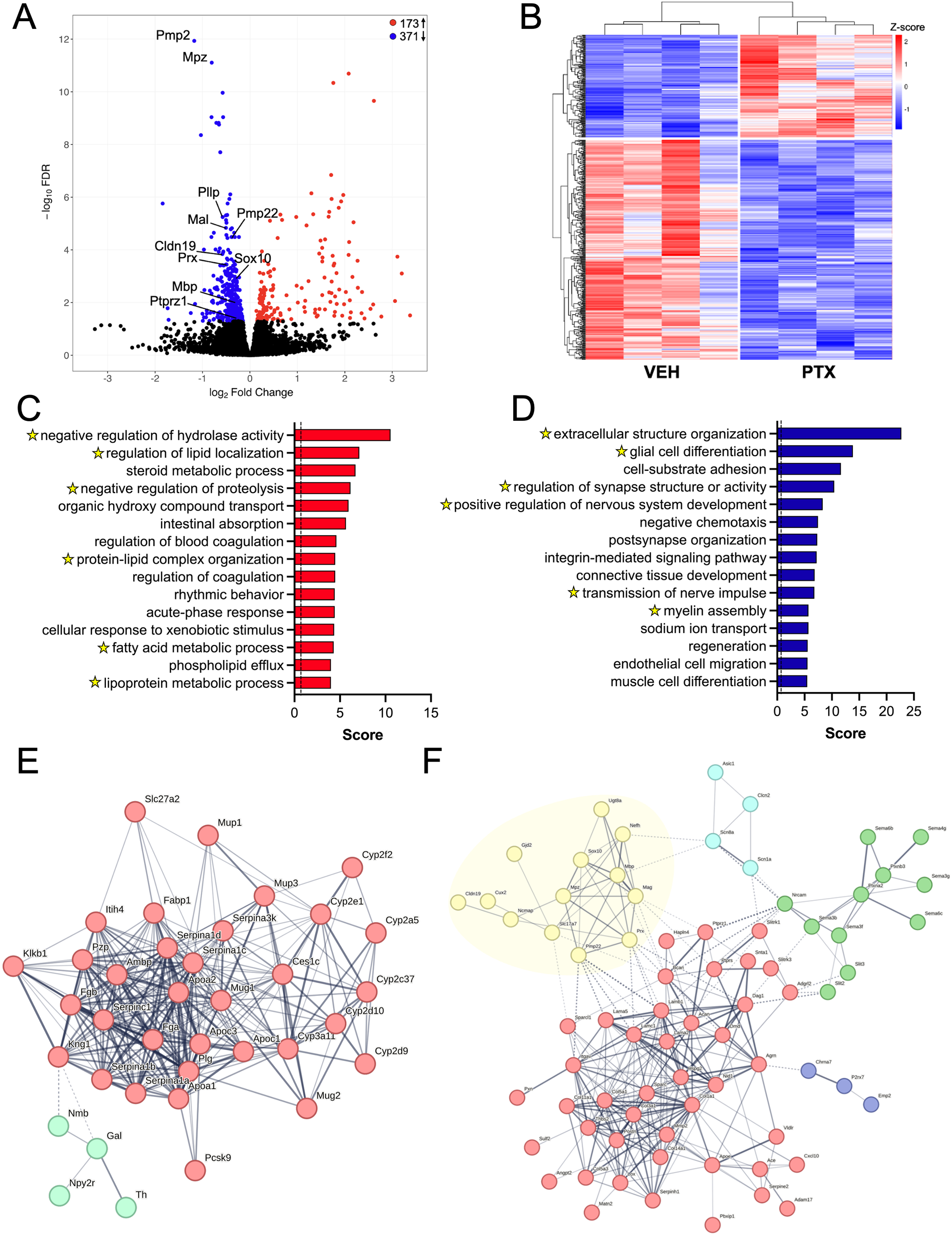
Consecutive PTX cycles cause cold hypersensitivity that persists longer than mechanical hypersensitivity. **(A)** Female mice received three cycles of PTX (4 mg/kg/dose) or vehicle where each cycle is made up of four doses (red arrows). Mechanical and cold sensitivity were assessed via the von Frey and acetone tests throughout the study. Tissue was harvested on day 145 (12 weeks after the third PTX cycle, black arrow). Animals were determined to have exclusive cold hypersensitivity days 120-145 (yellow shading). **(B)** Response (seconds) to acetone stimulation as a measure of cold sensitivity throughout three cycles of PTX. **(C)** Percent paw withdrawal frequency to 1.4g force filament as a measure of mechanical sensitivity throughout three cycles of PTX. **(D)** The average response (s) to acetone stimulation during exclusive cold hypersensitivity. **(E)** Area under the curve (AUC) of response (s) during exclusive cold hypersensitivity. Data is represented as mean ±SD. * *P* < 0.05 and **** *P* < 0.0001, (n=8/group).

### Behavioral assessments

Mechanical sensitivity was assessed to low-force (0.6 g) and high force (1.4 g) von Frey monofilaments following the low-high assessment method^36^ at baseline, twice weekly for 16 weeks, then once weekly for 4 weeks as previously described.^33^ Briefly, mechanical stimulation was applied to the plantar surface of the hind paw for 1 second. Stimulation was repeated for 10 trials with >5 minutes between each trial. The proportions of positive responses (i.e., abrupt lifting, shaking, extended lifting, and/or foot licking) observed from the 10 trials for the right and left paws were averaged and reported as percent paw withdrawal frequency (%PWF) for individual animals at each time point.

Cold sensitivity was assessed via the acetone test^37,38^ at baseline, twice weekly for 16 weeks, then once weekly for 4 weeks as previously described.^33^ Briefly, acetone was applied to the plantar surface of the hind paw via a syringe and the time spent responding (i.e., lifting, shaking, extended holding, and/or foot licking) to the acetone application was recorded for the subsequent 60 seconds. Acetone application was repeated five times with >5 minutes between each trial. To ensure accurate and unbiased assessments, mice were videorecorded during acetone testing and videos were scored blindly. The mean response time (seconds; right and left paws) across all five trials for each mouse at each time point were used for statistical analysis.

### RNA-sequencing and data processing

For each biological replicate, total RNA was extracted from right lumbar (L3-7) DRGs of one mouse using the QIAGEN RNeasy Mini Kit (QIAGEN, Germantown, MD) according to manufacturer’s instructions. RNA concentration was measured using the Qubit 4 Fluorometer with the Qubit RNA BR Assay Kit (Invitrogen, Carlsbad, CA) and integrity was assessed using RNA Nano Eukaryote chips on an Agilent 2100 Bioanalyzer (Agilent Technologies, Santa Clara, CA). The average concentration = 50 ng/uL and average RIN = 6.9. Strand-specific RNA sequencing libraries were constructed using 400 nanograms of total RNA and the NEBNext Ultra II RNA Library Prep Kit for Illumina with poly(A) selection (NEBNext Poly(A) mRNA Magnetic Isolation Module, New England Biolabs, Ipswich, MA) according to manufacturer’s instructions. Libraries (one mouse/sample) were barcoded using the recommended NEBNext Multiplex Oligos (New England Biolabs, Ipswich, MA). The size range and quality of each library was verified using High Sensitivity DNA chips on an Agilent 2100 Bioanalyzer (Agilent Technologies, Santa Clara, CA) and concentration was measured using the Qubit 4 Fluorometer with the Qubit DNA HS Assay Kit (Invitrogen, Carlsbad, CA). Libraries were diluted to 10nM and pooled in equimolar concentrations. Paired-end, 150 base pair sequencing was performed on a single lane on an Illumina NovaSeq X (Psomagen, Rockville, MD) to an average depth of 115.7 million reads per sample. Each group consisted of four independent biological replicates (1 mouse/replicate, n=4 mice/group) for a total of 8 libraries sequenced.

Sequencing reads were trimmed using Trim Galore^39^, assessed for quality by FastQC, and aligned to annotated RefSeq genes in the mouse reference genome (mm10) using HISAT2.^40^ A count matrix was generated using featureCounts^41^ and filtered to remove ribosomal RNA and genes with 0 reads for all samples. The remaining raw counts were imported into R for analysis. Default procedures of DESeq2^42^ were used to normalize, log transform, and analyze gene counts to identify genes that were differentially expressed between PTX and VEH groups. *P* values were corrected for multiple comparisons using Benjamini-Hochberg method^42^, and a false discovery rate (FDR) <0.05 was used to define differentially expressed transcripts between PTX and VEH groups. Differentially expressed genes (DEGs) underwent gene ontology (GO) pathway analysis to infer their biological processes using GOseq^43^ and results were simplified to their parent terms using the rrvgo package using a score cut off of 0.7.^44–46^ DEGs present in more than one of the top 10 GO parent terms were used for STRING analysis with minimum interaction score of 0.4 and were clustered by MCL clustering with an inflation parameter of _1.5._47,48

### Sciatic nerve processing, immunofluorescence, and analysis

Sciatic nerves were isolated and stored in 4% formaldehyde for 6 hours at room temperature before being washed with PBS and transferred to 20% glycerol for storage. Tissue was embedded in paraffin and cut into 4-μm-thick sections. Sections underwent antigen retrieval with 1X Citrate Buffer, pH 6.0 (Sigma-Aldrich, St. Louis, MO), blocking in 1X PBS containing 0.1% Tween-20 and 3% BSA for one hour at room temperature, and overnight incubation at 4°C with primary antibodies diluted in 1X PBS containing 0.1% Tween-20 and 1% BSA. The following primary antibodies were used: mouse anti-Mbp (Invitrogen, #MA5-15922, 1:2000), rabbit anti-Pgp9.5 (GeneTex, GTX109637, 1:500), rabbit anti-Iba1 (Fujifilm, 019-19741, 1:200), and rabbit anti-Egr2 (Abcam, ab245228, 1:500).Sections were washed three times with 1X PBS containing 0.1% Tween-20 prior to incubation with donkey anti-mouse Alexa Fluor-647 or donkey anti-rabbit 555-conjugated secondary antibodies (Jackson ImmunoResearch, 1:400) in blocking solution for one hour and washed as described above. Sections stained for Egr2 were counterstained with 0.1 µg/mL DAPI. Sections were coverslipped in ProLong Diamond Antifade Mountant (Invitrogen) and stored at 4°C.

Axon myelination was quantitatively assessed for the 1) area of myelin expression, 2) frequency of partial myelin sheaths, and 3) incidence of axonopathy in one section per animal from four animals per group (n=1 section/animal). Three non-overlapping images per section were acquired for each biological sample using a Zeiss LSM 880 confocal microscope and 100X oil immersion objective. Zeiss ZEN 2 software was used to acquire images and render z-stacks into maximum intensity projections for analysis. Images were acquired and scored blindly. The cell counter function in ImageJ 2.18.0 was first used to count axons by Pgp9.5 staining. The myelin sheath associated with each axon was assessed by Mbp staining and defined as full myelin sheaths (Mbp staining completely surrounding the axon), partial myelin sheaths (Mbp staining with gaps around the axon), or no myelin sheath (no Mbp staining associated with axon). Counts were converted into percentages by dividing each category by the total number of axons counted. Full myelin sheaths without a central axon were considered to be instances of axonopathy and results were converted into percentages by dividing the number of instances of axonopathy by the number of myelin sheaths assessed. Each image captured 100-300 axons and sheaths and the counts from each of the three images were averaged for each sample. The number of axons counted was divided by the area of the nerve examined to determine axons/µm^2^ as a measure of axonopathy and the ratio of Mbp/Pgp9.5 mean fluorescence intensity was used to assess the amount of Mbp.

Sections were quantitatively assessed for the number of Iba1-expressing cells. A single image per section was acquired for each biological sample using an Olympus BX51 microscope and 40X objective using MetaMorph software 7.7.0.0. The number of Iba1 positive regions that measured >3µm^2^ were counted using the analyze particles function in ImageJ 2.18.0 and expressed as the number of Iba1 expressing cells/µm^2^. Sections were also assessed for the number of Egr2-expressing cells. A single image per section was acquired for each biological sample using a Zeiss LSM 880 confocal microscope and 20x objective using Zeiss ZEN 2 software. The number of Egr2 positive regions >1µm^2^ that overlapped with a DAPI positive region were counted using the analyze particles function in ImageJ 2.18.0 and expressed as the number of Egr2 expressing cells/µm^2^. Images were acquired and scored by individuals blind to group assignment.

### ATAC-sequencing and data processing

Following dissection, left lumbar (L3-7) DRGs from one mouse were transferred directly to cold lysis buffer (0.32 M sucrose, 5 mM calcium chloride, 3 mM magnesium acetate, 10 mM Tris-hydrochloride, pH 8.0, 1 mM dithiothreitol, 0.1% TritonX-100, 0.1 mM EDTA). Nuclei were isolated by dounce homogenization of DRGs in lysis buffer followed by centrifugation at 350 x g at 4°C for five minutes. Nuclei were resuspended in 1X PBS and counted using a hemocytometer. Tn5 tagmentation was performed by OMNI_ATACseq as previously described^18,49^ with each 50 μL reaction containing 50,000 nuclei, 10mM Tris-HCl 7.6, 5mM MgCl2, 10% Dimethyl Formamide, 66% 1X PBS, 0.02% digitonin, 0.2% Tween-20, and 2.5 μL TDE1 enzyme (Illumina), and incubated at 37°C for 30 minutes. Tagmented DNA was purified with the QIAGEN MinElute PCR purification Kit (QIAGEN, Germantown, MD) according to manufacturer’s instructions. DNA fragments were amplified using Nextera Index adaptors, NEBNext High Fidelity 2X PCR Master Mix (New England Biolabs, Ipswich, MA) and 10 cycles of PCR. Libraries were then purified using the Zymo Clean and Concentrate-5 Kit (Zymo, Irvine, CA) according to manufacturer’s instructions. Size range and quality of each library was verified using High Sensitivity DNA chips on an Agilent 2100 Bioanalyzer (Agilent Technologies, Santa Clara, CA) and concentration was measured using the Qubit 4 Fluorometer with the Qubit DNA HS Assay Kit (Invitrogen, Carlsbad, CA). Libraries were diluted to 10nM and pooled in equimolar concentrations. Paired-end, 150 base pair sequencing was performed on a single lane on an Illumina NovaSeq X (Psomagen, Rockville, MD) to an average depth of 40 million reads per sample. Each group consisted of four independent biological replicates (1 animal/replicate) for a total of 8 libraries sequenced.

Sequencing reads were trimmed using Trim Galore^39^, assessed for quality by FastQC, and aligned to annotated RefSeq genes in the mouse reference genome (mm10) using Bowtie 2.^50^ Duplicate reads were removed using the *MarkDuplicates* function in Picard.^51^ Aligned reads were shifted 4 nucleotides upstream for the 5’ end and 5 nucleotides downstream for the 3’ end to account for potential artifacts from Tn5 binding. Bedtools^52^ *slop* function was used to extend the insertion sites by 75 base pairs up and downstream. Each sample was then downsampled to 52 million insertion sites to account for differences in sequencing depth. Peaks for each sample were called using the *callpeak* function in Model-based Analysis of ChIP-Seq (MACS2)^53^ using the flags -nomodel -extsize 150 -keep-dup all -call-summits. Bigwig files for each sample were generated for visualization in Integrative Genomics Viewer.^54^

Downsampled bam files were imported into DiffBind to define a consensus peakset which was subsequently used for differential accessibility analysis. Genomic regions called as peaks in at least 50% of biological replicates were included in the consensus peakset. The number of reads aligned to each region were counted and differential accessibility regions (DARs) between the PTX and VEH groups were determined using DiffBind at *P*<0.01.^55^ DARs were then annotated to the nearest annotated transcription start site using the *annotatePeaks.pl* function in Hypergeometric Optimization of Motif Enrichment (HOMER).^56^ The *findMotifsGenome.pl* function in HOMER was used to identify enrichment of transcription factor binding motifs within the differentially accessible regions.

### Cloning, transfection, and luciferase assays

Luciferase reporter constructs were constructed by cloning a candidate regulatory region into the pGL3 promoter vector (Promega; Madison, WI). Each region was inserted using standard restriction enzyme-based cloning techniques. Briefly, regions were obtained by PCR of mouse genomic DNA using NEBNext High Fidelity 2X PCR Master Mix (New England Biolabs, Ipswich, MA) with the 5’ end of the primers modified to contain BglII and MluI restriction sites respectively (Supplemental table 1). PCR products were digested and ligated into the BglII and MluI restriction enzyme sites of the pGL3 promoter vector. The ligated products were transformed into competent DH5α *E.coli* cells with ampicillin (100μg/mL) selection for plasmid-positive colonies. All constructs were verified by restriction enzyme digestion.

NIH3T3 cells (ATCC, Manassas, VA) were transfected with cloned pGL3 luciferase construct and pGL4.74 renilla expression vector for normalization using ViaFect Transfection Reagent (Promega; Madison, WI) as previously described.^18^ Luciferase and renilla expression were measured on the BioTek Synergy H1 microplate reader using the Dual-Glo Luciferase reporter assay system (Promega; Madison, WI) as previously described.^18^

### Statistical analysis

All data are represented as mean ± SD. The sample size for all experiments is noted in the figure legends. Data obtained for behavior, histological, and luciferase assays were analyzed by 2-way ANOVA followed by Tukey’s post hoc test for multiple comparisons, 1-way ANOVA followed by Dunnet’s post hoc test for multiple comparisons, or by Student’s t test with F test to compare variances for single comparisons. A p-value <0.05 was considered statistically significant. Statistical analysis was performed using GraphPad Prism version 10.4.1 for Windows (GraphPad Software, Boston, Massachusetts USA, www.graphpad.com).

## Results

### Consecutive PTX cycles produce persistent cold hypersensitivity that is longer lasting than mechanical hypersensitivity

Female mice received three consecutive cycles of PTX or vehicle to mimic the clinical dosing regimen, and mechanical and cold sensitivity were consistently monitored via von Frey and acetone testing (Fig. 1A). Following PTX, mice developed cold hypersensitivity [*F*(1, 14) =61.53, *P*<0.0001] that remained elevated throughout the duration of the study (Fig. 1B). Mice developed mechanical hypersensitivity [*F*(1, 14) =87.18, *P*<0.0001] after receiving PTX that returned to baseline levels following cycle 3 (Fig. 1C). Mechanical hypersensitivity to 1.4g and 0.6g stimulation (data not shown) returned to basal levels on day 114. However, PTX mice continued to exhibit increased response to acetone compared to VEH mice for at least an additional four weeks, until day 141 (Fig. 1B). During this period of exclusive cold hypersensitivity, the average response to acetone stimulation was increased in PTX mice compared to VEH mice [11.3±3.4 versus 7.2±2.0 seconds, t(3.001)=11.57, *P*=0.0115] (Fig. 1D). Similarly, the area under the curve of the response to acetone stimulation was increased in PTX mice compared to VEH mice [253.9±21.3 versus 129.6±15.8, t(4.684)=51.70, *P*<0.0001] (Fig. 1E).

### Persistent cold hypersensitivity is associated with the downregulation of genes involved in myelin maintenance

To understand the mechanisms that underlie PTX-induced cold hypersensitivity, we examined gene expression changes in the DRG of female mice during the period of exclusive cold hypersensitivity. We found 543 DEGs with 173 genes upregulated and 371 genes downregulated in PTX versus VEH mice (Fig. 2A-B). Upregulated DEGs are involved in biological processes of lipid and protein processing and localization (Fig. 2C) while the downregulated DEGs have roles in extracellular structure organization, nervous system development, transmission of nerve impulse, and myelin assembly (Fig. 2D). We found 83 DEGs that were shared in the top parent biological processes that we examined for relationships through STRING analysis and MCL clustering. Upregulated DEGs were clustered by complement and coagulation cascades (red) and neuropeptide binding (green) (Fig. 2E) and downregulated DEGs were clustered by extracellular matrix organization (red), myelination (yellow), semaphorin-plexin signaling pathway (green), sodium channel (blue), and bleb assembly (purple) (Fig. 2F).

**Figure 2.**
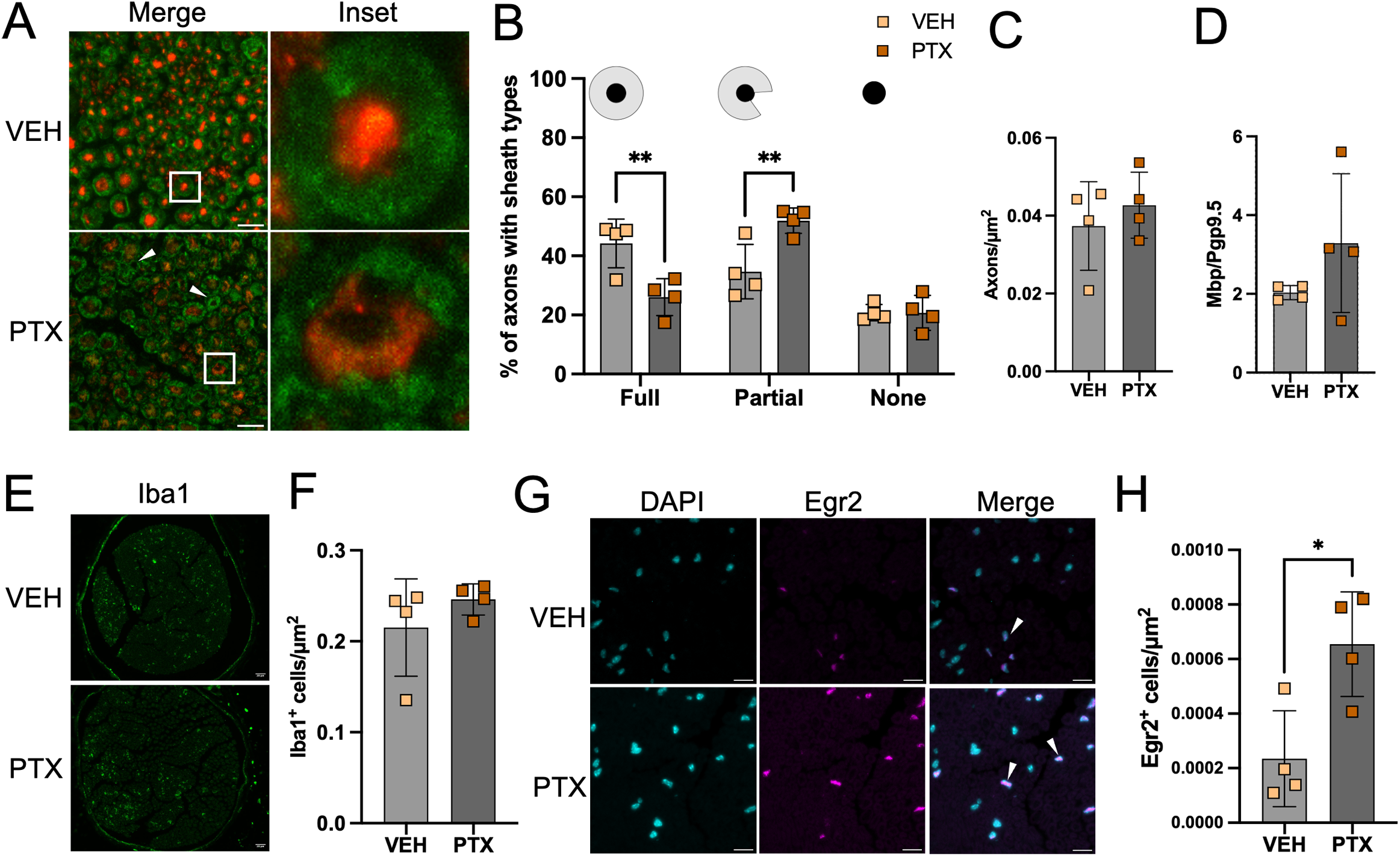
Genes involved in myelination are downregulated during persistent cold hypersensitivity. **(A)** Differentially expressed genes (DEGs) with *FDR* <0.05 in dorsal root ganglia (DRG) from female mice during exclusive cold hypersensitivity after PTX. **(B)** Replicate consistency of DEG expression within PTX and VEH groups. Gene ontology analysis of biological processes identified in upregulated **(C)** and downregulated **(D)** DEGs simplified to parent terms using rrvgo, pathways of interest containing DEGs present in multiple enriched GO parent terms are highlighted with yellow stars. **(E)** STRING analysis of upregulated DEGs present in multiple enriched GO parent terms clustered by complement and coagulation cascades (red) and neuropeptide binding (green). **(F)** STRING analysis of downregulated DEGs present in multiple enriched GO parent terms clustered by extracellular matrix organization (red), myelination (yellow), semaphoring-plexin signaling pathway (green), sodium channel (blue), and bleb assembly (purple). (n=4/group).

### Myelin sheaths in sciatic nerves are degraded during persistent cold hypersensitivity

Since genes involved in myelination were downregulated, we assessed morphology of sciatic nerves from PTX and VEH mice during this period of exclusive cold hypersensitivity (Fig. 3A). Compared to VEH animals, PTX animals had a lower percentage of axons with full myelin sheaths [26.1±6.3% versus 44.2±8.3%, t(3.952)=18, *P*=0.0028] and a higher proportion of axons with partial myelin sheaths [51.9±4.3% versus 34.7±9.2%, t(3.750)=18, *P*=0.0044] (Fig. 3B). We found no differences in the number of axons/µm^2^ or the ratio of Mbp/Pgp9.5 expression between PTX and VEH mice (Fig. 3C-D), however, the PTX animals had significantly higher variance in their Mbp/Pgp9.5 expression compared to the VEH group [*f*(95.08) = 3, *P*=0.0036] (Fig. 3D). We speculate that this variance reflects the uneven distribution of myelin debris in the PTX samples compared to the control ratio of 2:1 myelin to axon area. Because we found no mean change in Mbp expression despite increased fragmentation of the myelin sheaths, we hypothesize that PTX affects the ability of infiltrating macrophages to clear myelin debris in injured nerves. Therefore, we assessed the number of Iba1+ cells in the sciatic nerve from PTX and VEH mice (Fig. 3E) and found no difference in the number or activation status of macrophages (Fig. 3F). We also assessed the expression of the pro-myelinating transcription factor Egr2 (Fig. 3G) by the colocalization of Egr2 and DAPI signals. We found that PTX mice had more Egr2+ cells in the sciatic nerve compared to VEH mice (Fig. 3H).

**Figure 3.**
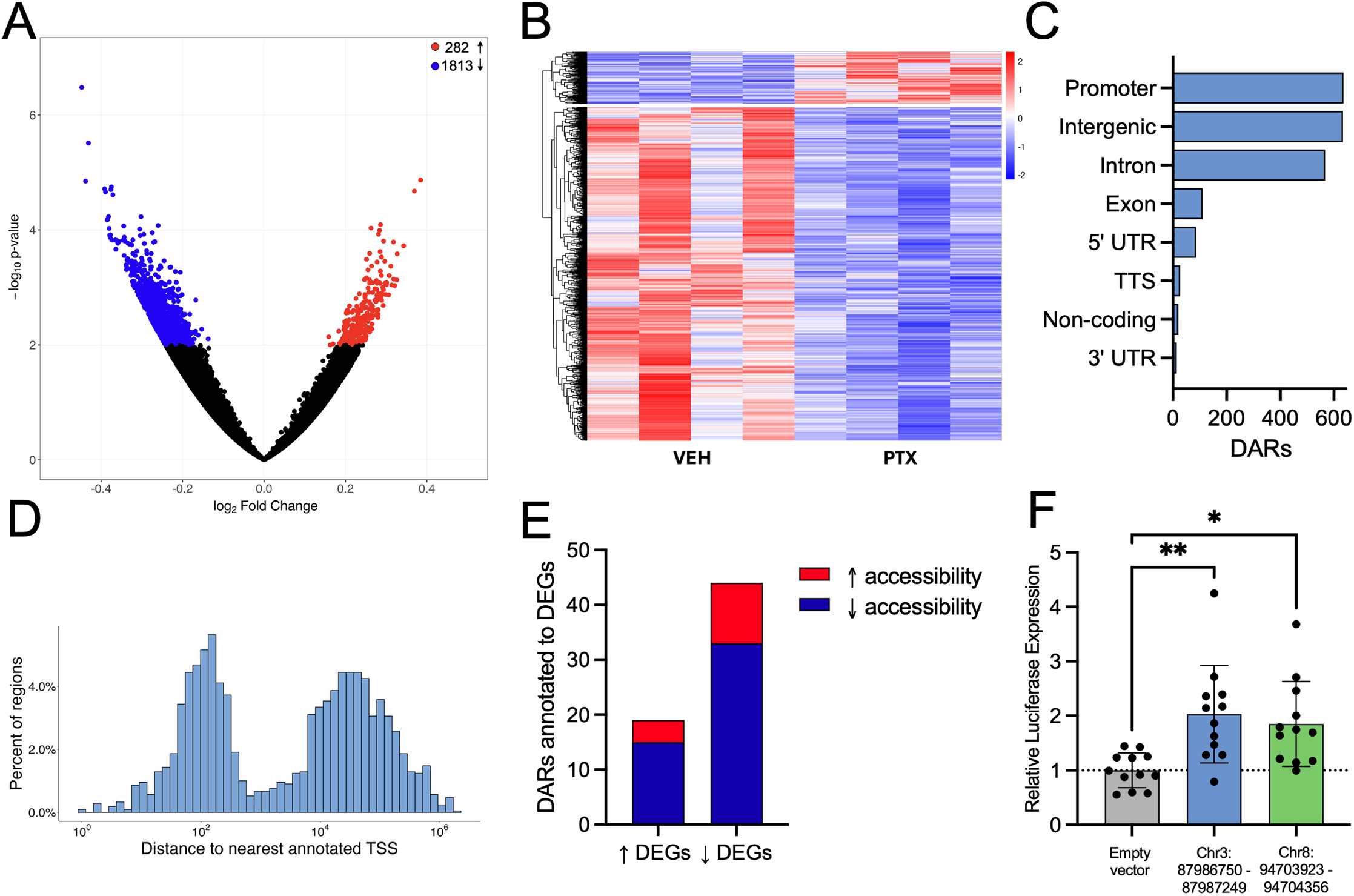
Myelin sheaths in the sciatic nerve are degraded during persistent cold hypersensitivity. **(A)** Representative images of sciatic nerves stained for Pgp9.5 and Mbp shows degraded myelin sheaths and axonopathy (arrow heads) in animals who received PTX. Scale bar = 10µm **(B)** The percentage of axons counted with full, partial, or no myelin sheath. **(C)** The number of axons/µm^2^. **(D)** The ratio of MFI of Mbp/Pgp9.5. **(E)** Representative images of sciatic nerves stained for Iba1. Scale bar=20µm **(F)** The number of Iba1+ cells counted per area. **(G)** Representative images of sciatic nerves stained for Egr2 and DAPI. Scale bar=10µm **(H)** The number of Egr2+ cells counted per area. Data is represented as mean ±SD. * *P*<0.05, ** *P* < 0.01, (n=4/group).

### Cold hypersensitivity is associated with long-lasting changes in chromatin accessibility at regions annotated to genes involved in myelination

To identify regulatory changes that underlie differential gene expression during persistent cold hypersensitivity, we used ATAC-sequencing to examine the chromatin accessibility in the DRGs of the same female mice used for RNA-sequencing. We identified 2095 DARs with 282 regions showing increased accessibility and 1813 regions showing decreased accessibility in PTX mice compared to VEH mice (Fig. 4A-B). We found that 30.4% of DARs were associated with annotated promoters and 57.4% were in intergenic and intronic regions (Fig. 4C). The genomic distance of these DARs from the nearest annotated transcription start site exhibited the expected bimodal distribution with peaks representing promoters (<1000bp) and distal regulatory regions (10-100kb) (Fig. 4D). 63 DARs were annotated to DEGs, we identified 36 of these DARs with concordant changes in gene expression, suggesting potential enhancer activity (Fig. 4E; Supplemental table 2). The DARs Chr3:87986750-87987249 and Chr8:94703923-94704356 were significantly less accessible in PTX DRGs compared to VEH DRGs (Supplemental Figure 1) and were annotated to the *Bcan* and *Pllp* genes, which were both downregulated in PTX mice. These DARs produced significantly greater luciferase expression compared to an empty vector control [*F*(2,33)=7.219, p=0.0025], indicating that both Chr3:87986750-87987249 [2.030±0.896 versus 1.000±0.320, *q*(3.552)=33, *P*=0.0023] and Chr8:94703955-94704650 [1.854±0.779 versus 1.000±0.320, *q*(2.944)=33, *P*=0.0112] possesses transcription enhancer abilities (Fig. 4F).

**Figure 4.**
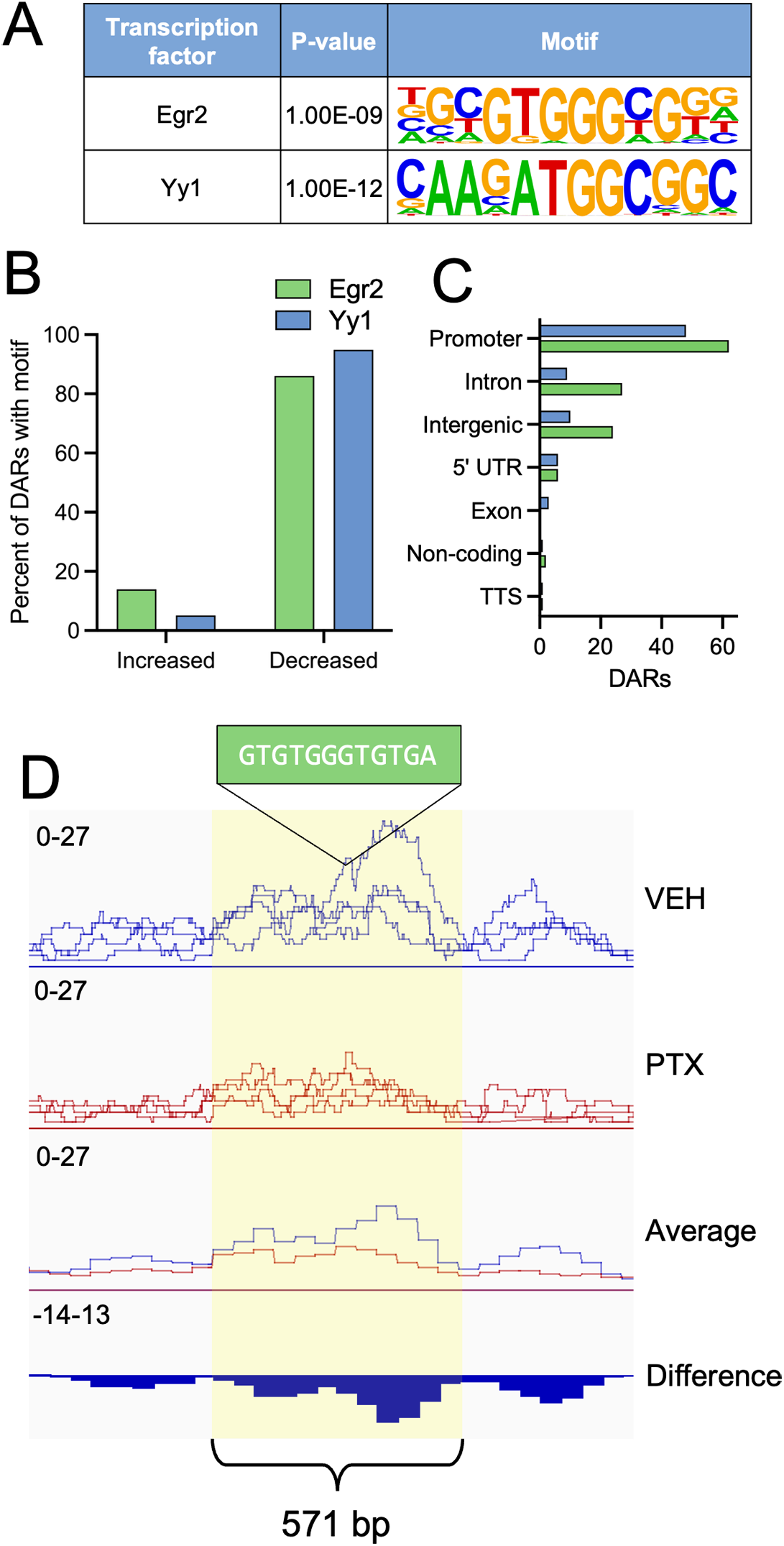
Differential chromatin accessibility is linked to downregulated myelin genes during cold hypersensitivity. **(A)** Differentially accessible regions (DARs) with *P* < 0.001 in DRG of female mice during exclusive cold hypersensitivity after PTX. **(B)** Replicate consistency of DAR accessibility within PTX and VEH groups. **(C)** Location of DARs across genomic features. **(D)** Distribution of DARs across distance to nearest annotated transcription start site. **(E)** DAR accessibility relative to the annotated DEG’s expression. **(F)** Accessibility of Chr3:87986750-87987249. **(G)** Relative luciferase expression in NIH3T3 cells transfected with pGL3 promoter vector or pGL3 promoter vector containing Chr3:87986750-87987249 and Chr8:94703923-94704356. (n=4/group).

### Myelin-regulating transcription factor Egr2 has binding motifs in DARs associated with DEGs

As our ATAC-seq identified DARs with potential regulatory function, we next sought to identify candidate transcription factors that may regulate genes that contribute to persistent cold hypersensitivity. Binding motifs for Egr2 and Yy1 were significantly enriched in DARs during the period of exclusive cold hypersensitivity (Fig. 5A). Interestingly, of the 200 DARs containing Egr2 and Yy1 binding motifs, 90.5% had decreased accessibility in PTX animals (Fig. 5B) and 55% were in promoter regions (Fig. 5C). For example, Chr18:82525917-82526487, with decreased accessibility in PTX DRGs, contains the Egr2 binding motif and is located 28kb from *Mbp*, the major constituent of the myelin sheath, which was downregulated in PTX mice during cold hypersensitivity (Fig. 5D).

**Figure 5.**
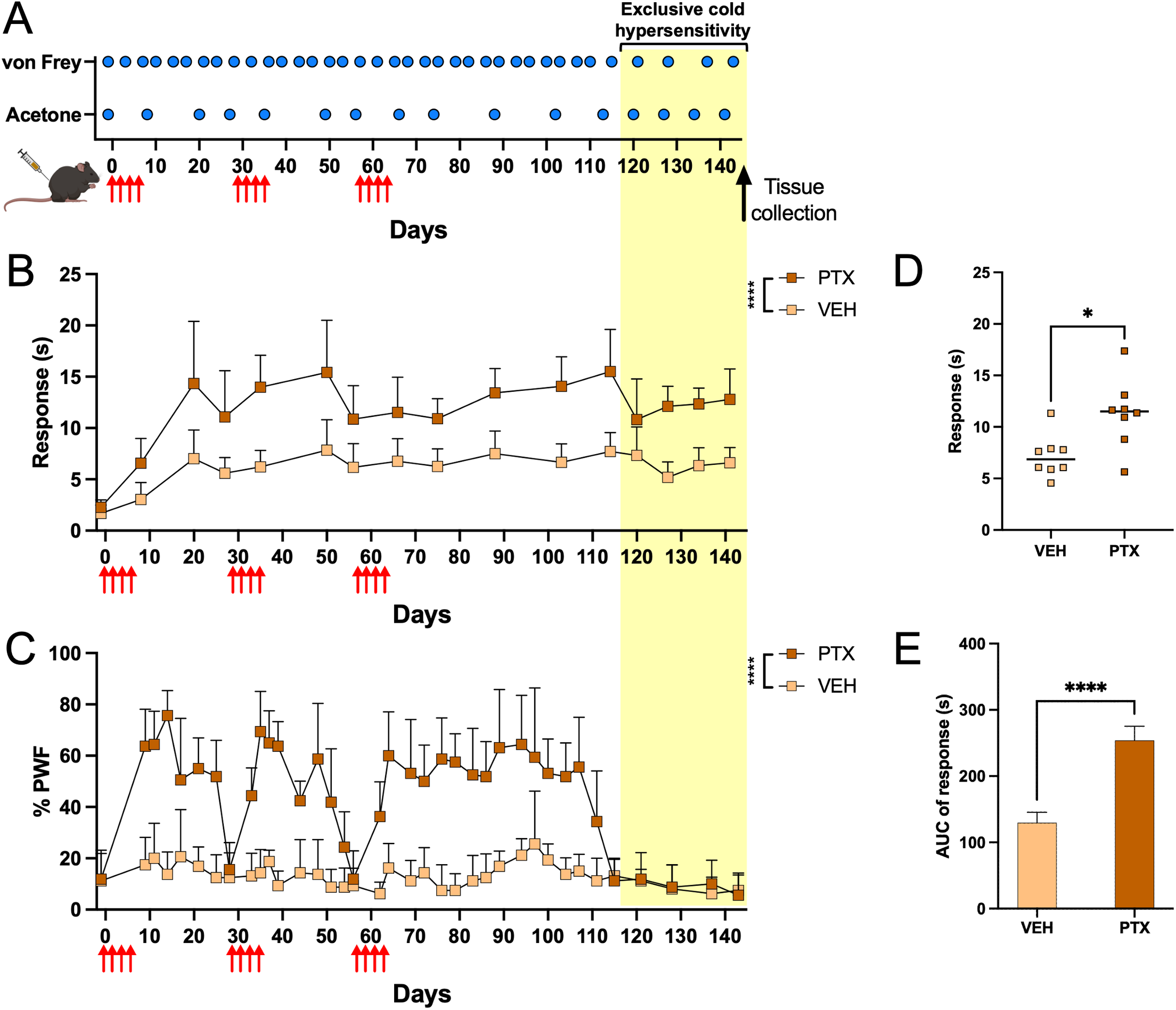
Motifs for myelin-regulating transcription factors are enriched in differentially accessible chromatin regions. **(A)** Motif analysis with HOMER revealed significant enrichment of EGR2 and YY1 binding motifs in DARs. **(B)** Accessibility of DARs containing Egr2 and Yy1 binding motifs. **(C)** Location of DARs containing Egr2 and Yy1 binding motifs across genomic features. **(D)** Chromatin accessibility of Chr18:82525917-82526487, intronic region of *Mbp* gene containing Egr2 binding motif at 243-255bp of the 571bp region.

## Discussion

Our study appears to be the first to examine pathological cold hypersensitivity in the absence of mechanical hypersensitivity. We found that PTX induced cold hypersensitivity that persisted for 12 weeks after the third PTX cycle and at least four weeks after the resolution of mechanical hypersensitivity. During this period of persistent exclusive cold hypersensitivity, DRGs from mice that received PTX showed downregulation of genes involved in axon myelination and changes in chromatin accessibility in putative cis-regulatory regions that contain binding motifs for the pro-myelinating transcription factors Egr2 and Yy1 and the sciatic nerves of these mice had increased fragmentation of myelin sheaths despite increased expression of Egr2. These findings indicate a coordinated signature of decreased chromatin accessibility at Egr2 and Yy1 motifs, decreased myelin gene expression, and degraded axon myelination during exclusive PTX-induced cold hypersensitivity and nominates Egr2-regulated myelin programs as candidate drivers of cold allodynia.

Despite advancements in our understanding of the molecular mechanisms underlying cold nociception, the drivers of the development and persistence of pathological cold pain remain unclear.^1,57^ An improved understanding of these mechanisms is essential for the development of preventative and therapeutic strategies for patients living with chronic cold pain.^1,5,58^ We previously demonstrated that PTX-induced cold and mechanical hypersensitivities have unique trajectories of development and resolution.^33^ In mice that receive three consecutive cycles of PTX, mechanical hypersensitivity returns to baseline level within 30 days after cycles 1 and 2 and within 60 days following the third PTX cycle.^33^ However, cold hypersensitivity does not return to baseline levels between PTX cycles, consistent with clinical reports that cold allodynia can be irreversible between cycles.^33^ Further, cold hypersensitivity that persists for at least 12 weeks after the third PTX cycle, four weeks longer than mechanical hypersensitivity. These findings agree with previous work that has demonstrated that PTX and other chemotherapeutics, induces cold hypersensitivity.^35,59,60^ Previous work shows cold sensitivity returning to baseline levels 22 days after a single cycle of PTX^35^ while other work shows it is maintained through day 30 after PTX,^59^ or even 10 weeks after oxaliplatin.^60^ Unfortunately, rodent models using chemotherapy to induce pathological cold pain also induce mechanical hypersensitivity,^61–64^ however, we are able to examine changes associated with cold hypersensitivity without the presence of confounding mechanical hypersensitivity. While all mice used in our study were treated and assessed in parallel as a single cohort, due to the length of the model, mice were well advanced in adulthood during our assessments of exclusive cold hypersensitivity. Nonetheless, this translational model mimics the clinical scenario of long-lasting cold allodynia and enables us to study the mechanisms that underlie pathological cold pain.

Cold is detected by the primary sensory neurons in dorsal root ganglia (DRG) via temperature-sensitive ion channels expressed on thin myelinated (Aδ) and unmyelinated (C) fibers that innervate the skin.^10,65–67^ Changes in gene transcription in the DRG alter peripheral nerve function and pain signaling and may underlie chronic hypersensitivity conditions.^64,68,69^ Our current understanding of the mechanisms that contribute to cold pain stems from work utilizing CIPN models,^24,25,28,29^ nerve injury models,^26,30^ and *in vivo* and *ex vivo* electrophysiology studies.^27,31,70,71^ Previous work supports the role of Trpm8 in the transduction of cold stimuli and shows increased *Trpm8* expression in the DRG during early onset cold hypersensitivity.^24–27^ In the present study, we found *Trpm8* expression to be decreased, further investigation is needed to understand the role of Trpm8 and its expression dynamics in cold hypersensitivity development and chronification. Previous studies also associate cold hypersensitivity with reduced expression of voltage-gated potassium channels, which depolarize the neuronal membrane and prevent continual action-potential propagation.^28–32^ We observed reduced expression of multiple potassium channels in the DRG during cold hypersensitivity, including *Kcnc1* and *Kcnc3*, suggesting that an impairment in ion handling in neurons may underlie pathological cold pain.^28–32^ ^24–27^ ^28–31^We also found that *Kcnip2* and *Kcnq1* expressions were increased. Genetic ablation of these channels reduces cold sensitivity, indicating that their increased expression may contribute to cold allodynia.^70,71^ Importantly, studies that utilized models of CIPN and nerve injury to examine mechanisms of cold pain identified changes associated with the presence of both cold and mechanical hypersensitivity.^26,28–30^ This limitation prevents the disaggregation of modality-specific findings to identify the mechanisms that specifically underlie cold hypersensitivity.

In the present study, we examined transcriptomic changes in the DRG four weeks after the resolution of mechanical hypersensitivity, when only cold hypersensitivity was present. We observe decreased expression of genes that encode for critical components of the myelin sheath, such as *Mbp* and *Mpz*, and components necessary for proper myelin sheath formation and function, including *Sox10* and *Pmp2*.^72,73^ Genes involved in the semaphorin-plexin signaling pathway, which is involved in cell morphology and motility, were also downregulated.^74^ Interestingly, decreased semaphorin-plexin signaling delays oligodendrocyte maturation thereby reducing myelination in the central nervous system, supporting the contribution of this pathway to decreased myelination during cold hypersensitivity.^75^ Demyelination leads to chronic pain and cold allodynia due to dysregulated ion handling and nerve conductance.^12,23,76,77^ Our findings implicate maladaptive transcriptional changes in the expression of genes encoding myelin components and potassium channels as primary contributors to cold hypersensitivity. However, our use of bulk RNA-sequencing of the DRG prevents us from assessing cell type composition and cell-type specific transcriptional changes between groups. Future single cell transcriptomic studies could clarify how the downregulation of myelin-associated genes in Schwann cells is linked to differential expression and hyperexcitability in specific neuronal subtypes.

Transcription factor binding at cis-regulatory elements (CREs) coordinates the timing, extent, and context of gene expression and therefore has significant influence in the regulation of cell signaling mechanisms.^78^ Chromatin accessibility, and therefore transcription factor binding, at CREs is influenced by environmental stimuli.^18,20^ Previous work has identified changes in the accessibility of putative CREs in the context of pain; however, these changes have not been examined during exclusive cold hypersensitivity.^18,20^ In the present study, we found that PTX-induced cold hypersensitivity is associated with changes in chromatin accessibility. Binding motifs for the transcription factors Egr2 and Yy1, which promote gene transcription necessary for Schwann cell development and myelinating function,^73,78,79^ were enriched in DARs with decreased accessibility. We also observed decreased accessibility at DARs proximal to downregulated genes involved in myelin sheath formation and maintenance (i.e., *Mbp, Pllp,* and *Prx*). Our findings indicate that consecutive PTX cycles may promote decreased chromatin accessibility at Egr2 and Yy1 binding sites which reduces the transcription of genes necessary for the maintenance of myelinated axons, thereby contributing to cold hypersensitivity. Future work is needed to determine the cell type specific changes in chromatin accessibility at these loci, examine what binding sites are necessary for Egr2 and Yy1 function to allow for proper myelin gene expression, and how this transcription factor function may be modulated to alleviate cold hypersensitivity.

Fragmented myelin sheaths are associated with the redistribution of sodium channels away from Nodes of Ranvier and across the axonal surface while potassium channels decrease in the juxtaparanode and redistribute to the paranode.^80–82^ This reallocation of ion channels alters extracellular ion concentrations, prevents potassium channels from repolarizing the neuronal membrane to oppose action potential initiation, and promotes the spontaneous neuronal activity present in cold allodynia.^12,23,77^ In the present study, the sciatic nerve of PTX mice contained a greater proportion of axons with fragmented myelin sheaths compared to VEH mice, which supports an association between degraded myelin and exclusive cold hypersensitivity, and is consistent with prior CIPN studies reporting myelin degradation.^83–86^ However, myelinated A fibers transmit mechanical sensation,^87^ and myelin damage has been associated with mechanical hypersensitivity^88,89^ which appears contrary to our findings. Nevertheless, myelin damage can induce neuronal hyperexcitability even in unmyelinated C fibers.^90^

Myelin debris is degraded and phagocytosed by activated macrophages within ten days after nerve injury.^91,92^ This phagocytosis leads to Schwann cells transitioning into an active, pro-myelinating state to remyelinate the axon.^91,93,94^ We found fragmented myelin and a lack of activated macrophages in the sciatic nerve 82 days after PTX exposure, which suggests that these mechanisms of myelin clearance and remyelination are impaired. However, we observed an increase in Egr2+ cells in the sciatic nerves of animals with cold hypersensitivity, indicating that these Schwann cells were in an a promyelinating state in response to the presence of damaged myelin sheaths.^73,95,96^ Interestingly, this upregulation did not lead to remyelination of the axons, likely due to the decreased chromatin accessibility at regulatory regions necessary for Egr2 binding and subsequent transcription of myelinating genes.^73,97^ Egr2 is known to be required for proper nerve myelination^98^ but previous work has shown that increased Egr2 expression may not lead to remyelination in an *in vivo* nerve repair model,^99^ possibly due to limited transcription factor function. Our findings indicate that although the Schwann cells are in a pro-myelinating state during cold hypersensitivity, decreased chromatin accessibility following repeated PTX cycles prevents DNA binding, myelin gene transcription, and perpetuates myelin damage. Further examination of microglia activation and stimulation of Egr2 expression following repeated cycles of PTX is needed to understand the mechanism leading to persistent myelin degradation without proper remyelination.

PTX upregulates the transcription of pro-inflammatory genes in the DRG,^23,100^ which are necessary for the activation of macrophages and subsequent myelin debris clearance.^80^ In addition, we do not observe an increase in the expression of pro-inflammatory markers in this work, supporting the hypothesis that PTX-induced inflammation contributes to mechanical, rather than cold, pain.^23,101^ We hypothesize that the PTX-induced pro-inflammatory environment promotes mechanical hypersensitivity and that this inflammation is attenuated following the third PTX cycle, which prevents debris clearance and myelin repair mechanism activation, therefore maintaining cold hypersensitivity. Longitudinal examination of the myelin sheath during both cold and mechanical hypersensitivity throughout three cycles of PTX is needed to understand how the resolution of mechanical hypersensitivity corresponds to myelin sheath damage.

To our knowledge, this is the first study to examine the mechanisms contributing specifically to PTX-induced cold hypersensitivity. Our findings support an association between persistent cold hypersensitivity and changes in chromatin accessibility, the downregulation of myelinating genes, the prevention of axon remyelination, and the impairment of sensory neuron function due to degraded myelin sheaths. Further, our findings implicate a breakdown in normal macrophage-Schwann cell signaling in response to myelin damage, leading to the persistent myelin degradation observed during cold hypersensitivity. Our work highlights the importance of assessments of the specific mechanisms underlying cold hypersensitivity to develop targeted therapeutics and improve the quality of life of chronic pain patients.

## Data availability

Sequencing data will be made available via NCBI GEO. All data are available upon reasonable request to the corresponding author.

## Acknowledgements

We thank Jeff Kamykowski at the University of Arkansas for Medical Sciences Digital Microscopy Core for his help and expertise in image acquisition and Brandon Fowlkes, Craig Everett, Mike Miller, and Gary Pride at the Arkansas Children’s Research Institute for their help in caring for animals used in this study.

## Funding

This study was funded by the National Institutes of Health (P20GM121293), the Arkansas Children’s Research Institute, the University of Arkansas for Medical Science Winthrop P. Rockefeller Cancer Institute, and a Seeds of Science research grant. The funders had no role in the study design, data collection and analysis, decision to publish, or preparation of the manuscript.

## Competing interests

The author reports no competing interests.

## Supplementary material

Supplementary material is available online

## Notes

### Competing Interest Statement

The authors have declared no competing interest.

### Summary of Updates

Differential accessibilikty results and motif analysis to indicate a potential role of Egr2

