## Supplemental information for "Degraded myelin is associated with cold hypersensitivity in paclitaxel-induced peripheral neuropathy" (from another version of this preprint: version 1, 2026-06-02)

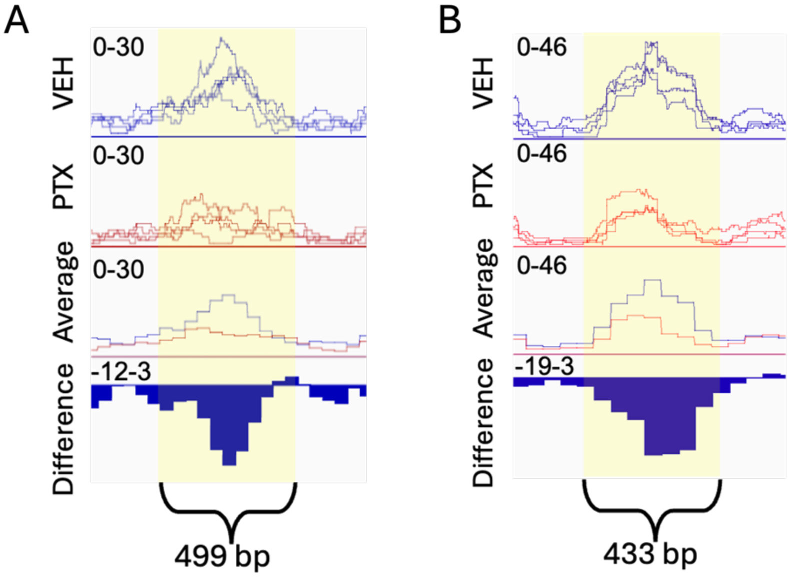


**Supplemental Figure 1 Differential chromatin accessibility at regions of interest during persistent cold hypersensitivity.** Differentially chromatin accessibility at **(A)** Chr3:87986750-87987249 and **(B)** Chr8:94703923-94704356. (n=4/group).

| Region of interest | Forward primer | Reverse primer |
|---|---|---|
| Chr3:87986750-87987249 | TGGAAGATCTTTGTTCTTGTCGCCTGGCTTTT | GCATACGCGTCCTGAGAGTGCCTCCGTTCC |
| Chr8:94703923-94704356 | TGGAAGATCTCCCCGCTCCTGGAATTCTTTTT | GCATACGCGTGGACCTGATTGCCACGTACC |

**Supplemental Table 1 Primers for cloning.** Primers used for restriction enzyme cloning of regions Chr3:87986750-87987249 and Chr8:94703923-94704356.

| Gene Name | Entrez ID | log2FC of transcript | p value of transcript | Region | log2FC of region | p value of region | Region width | Annotation |
|---|---|---|---|---|---|---|---|---|
| 5730507C01Rik | 236366 | 1.2346 | 0.001534 | chr12:18695785-18696079 | 0.2205 | 0.005152 | 295 | Intergenic |
| Agrn | 11603 | -0.3271 | 0.001364 | chr4:156197062-156198249 | -0.2492 | 0.005931 | 1188 | promoter-TSS |
| Bach1 | 12013 | -0.3042 | 0.026710 | chr16:87801299-87801468 | -0.2227 | 0.002134 | 170 | Intergenic |
| Bcan | 12032 | -0.2476 | 0.039889 | chr3:87986750-87987249 | -0.2693 | 0.003349 | 500 | TTS |
| Bcan | 12032 | -0.2476 | 0.039889 | chr3:87994224-87994763 | -0.2215 | 0.006327 | 540 | intron |
| Chdh | 218865 | -0.4947 | 0.000009 | chr14:30133107-30133655 | -0.2222 | 0.007840 | 549 | intron |
| Cldn19 | 242653 | -0.5809 | 0.000144 | chr4:119247507-119247928 | -0.2458 | 0.006303 | 422 | intron |
| Clic4 | 29876 | -0.3078 | 0.005280 | chr4:135272242-135273236 | -0.2798 | 0.001167 | 995 | promoter-TSS |
| Cotl1 | 72042 | -0.2946 | 0.048772 | chr8:119840095-119841087 | -0.2516 | 0.006363 | 993 | promoter-TSS |
| Csmd2 | 329942 | -0.3096 | 0.011208 | chr4:128087134-128087558 | -0.2523 | 0.003629 | 425 | intron |
| Ddit4 | 74747 | -0.4405 | 0.002456 | chr10:59943527-59944016 | -0.2450 | 0.003928 | 490 | Intergenic |
| Elovl1 | 54325 | -0.2919 | 0.021459 | chr4:118427734-118428603 | -0.2999 | 0.001040 | 870 | promoter-TSS |
| Gprc5b | 64297 | -0.3562 | 0.004889 | chr7:119023648-119023876 | -0.2063 | 0.008901 | 229 | Intergenic |
| Iqsec1 | 232227 | -0.3093 | 0.048573 | chr6:90716058-90716741 | -0.2364 | 0.008779 | 684 | 5' UTR |
| Itga6 | 16403 | -0.3436 | 0.000569 | chr2:71786576-71787918 | -0.3115 | 0.001180 | 1343 | exon |
| Itgav | 16410 | -0.2843 | 0.005529 | chr2:83754046-83754544 | -0.2561 | 0.002580 | 499 | intron |
| Kcnc3 | 16504 | -0.5548 | 0.000391 | chr7:44590512-44591432 | -0.3083 | 0.000848 | 921 | exon |
| Kcnip4 | 80334 | 0.2432 | 0.024940 | chr5:49583707-49583971 | 0.1868 | 0.009900 | 265 | Intergenic |
| Mbp | 17196 | -0.3928 | 0.012020 | chr18:82525917-82526487 | -0.2307 | 0.008557 | 571 | intron |
| Mug1 | 17836 | 2.3526 | 0.025404 | chr6:121780681-121780973 | 0.1848 | 0.009458 | 293 | Intergenic |
| Papln | 170721 | -0.8086 | 0.004546 | chr12:83792562-83792845 | -0.2393 | 0.004562 | 284 | TTS |
| Parp3 | 235587 | -0.3549 | 0.000396 | chr9:106474503-106474698 | -0.2552 | 0.004459 | 196 | intron |
| Per1 | 18626 | -0.4836 | 0.036892 | chr11:69096782-69097358 | -0.2505 | 0.002781 | 577 | Intergenic |
| Per1 | 18626 | -0.4836 | 0.036892 | chr11:69094677-69095463 | -0.2480 | 0.002598 | 787 | Intergenic |
| Per1 | 18626 | -0.4836 | 0.036892 | chr11:69098675-69099033 | -0.2400 | 0.005018 | 359 | promoter-TSS |
| Plekhg2 | 101497 | -0.4135 | 0.000988 | chr7:28375502-28375862 | -0.2439 | 0.001153 | 361 | Intergenic |
| Pllp | 67801 | -0.5750 | 0.000006 | chr8:94703955-94704650 | -0.2282 | 0.009023 | 696 | Intergenic |
| Prex2 | 109294 | -0.3745 | 0.023590 | chr1:11144536-11144952 | -0.1971 | 0.004511 | 417 | intron |
| Prx | 19153 | -0.4932 | 0.000360 | chr7:27507736-27508583 | -0.2878 | 0.001827 | 848 | intron |
| Relt | 320100 | -0.4846 | 0.047572 | chr7:100862738-100863939 | -0.2698 | 0.004297 | 848 | promoter-TSS |
| Sema6b | 20359 | -0.5218 | 0.008366 | chr17:56133848-56134242 | -0.2566 | 0.005981 | 395 | promoter-TSS |
| Septin9 | 53860 | -0.2565 | 0.018349 | chr11:117318422-117319018 | -0.2687 | 0.004603 | 597 | intron |
| Serpine2 | 20720 | -0.2541 | 0.023590 | chr1:80030816-80031601 | -0.2201 | 0.008779 | 786 | Intergenic |
| Sez6 | 20370 | -0.2328 | 0.028089 | chr11:77930748-77931223 | -0.3063 | 0.000990 | 476 | 5' UTR |
| Slitrk1 | 76965 | -0.3466 | 0.008437 | chr14:108650093-108650367 | -0.2667 | 0.000600 | 275 | Intergenic |
| Tafa4 | 320701 | 0.2819 | 0.037206 | chr6:96622700-96623311 | 0.2480 | 0.006358 | 612 | intron |

**Supplemental Table 2 DARs annotated to myelin-related DEGs.** 36 DARs annotated to DEGs related to myelin processes.
